# Study on gene knockout mice and human mutant individual reveals absence of CEP78 causes photoreceptor and sperm flagella impairments

**DOI:** 10.1101/2022.01.25.477668

**Authors:** Tianyu Zhu, Yuxin Zhang, Xunlun Sheng, Xiangzheng Zhang, Yu Chen, Yueshuai Guo, Yaling Qi, Yichen Zhao, Qi Zhou, Xue Chen, Xuejiang Guo, Chen Zhao

**Author notes:** These authors contributed equally to this work (Tianyu Zhu, Yuxin Zhang, Xunlun Sheng). Correspondence: Chen Zhao or Xuejiang Guo or Xue Chen.

## Abstract

**Background:** Cone-rod dystrophy (CRD) is a genetically inherited retinal disease characterized by photoreceptor degeneration. In some rare cases, CRD and hearing loss can be associated with male fertility, while the underlying mechanism is not well known.

**Methods:** Using CRISPR/Cas9 system, we generated *Cep78^-/-^* mice. And electroretinogram (ERG), immunofluorescence staining and transmission electron microscopy (TEM) were used to analyze visual function and photoreceptor ciliary structure changes in *Cep78^-/-^* mouse. HE/PAS staining, scanning-electron microscopy (SEM) were conducted to *Cep78^-/-^* mice and human CRD patient with CEP78 protein loss to illustrate male infertility and multiple morphological abnormalities of the sperm flagella (MMAF) caused by CEP78 deficiency. TEM and immunofluorescence staining were performed to characterize morphological and molecular changes of sperm flagella microtubule arrangement, centriole development and spermatid head shaping in *Cep78^-/-^* mice. Mass-spectrometry analyses were conducted to identify protein abnormalities after *Cep78* deletion and Cep78 interacting proteins in spermiogenesis. Co-immunoprecipitation was used to show the Cep78-Ift20-Ttc21a trimer. The role of Cep78-Ift20-Ttc21a trimer in cilliogenesis and centriole elongation was assessed by cilia induction assay.

**Results:** *Cep78* knockout mice exhibited impaired function and morphology of photoreceptors, typified by reduced electroretinogram amplitudes, disrupted translocation of cone arrestin, attenuated and disorganized photoreceptor outer segments (OS) disks and widen OS bases, as well as interrupted cilia elongations and structures. *Cep78* deletion also caused male infertility and MMAF, with disordered “9 + 2” structure and triplet microtubules in sperm flagella. CEP78 forms a trimer with intraflagellar transport (IFT) proteins IFT20 and TTC21A essential for sperm flagella formation, is essential for their interaction and stability, and recruits IFT20 to centrosome. Insufficiency of any component in the trimer causes centriole elongation and cilia shortening. Additionally, absence of CEP78 protein in human leaded to similar phenotypes in vision and MMAF as *Cep78^-/-^* mice.

**Conclusions:** We found *CEP78* as the causative gene of CRD with MMAF in human and mouse. Cep78 forms a trimer with Ift20 and Ttc21a, and regulate the interaction, stability and localization of the trimer proteins, which regulate cilliogenesis, centriole length, and sperm flagella formation.

**Funding:** This work was supported by the National Key R&D Program (2021YFC2700200 to X.G); National Natural Science Foundation of China (82020108006, 81730025 to C.Z, 81971439, 81771641 to X.G, 82070974 to X.C, 82060183 to X.S); Shanghai Outstanding Academic Leaders (2017BR013 to C.Z); and Six Talent Peaks Project in Jiangsu Province (YY-019 to X.G). The funders had no role in study design, data collection and analysis, decision to publish, or preparation of the manuscript.

## Introduction

Cone-rod dystrophy (CRD) is a genetically inherited retinal disease characterized by photoreceptor degeneration[1]. Many CRD pathogenic genes are associated with connecting cilia in photoreceptor, a type of non-motile cilia. CRD usually exists with combination of immotile cilia defects in other systems. CEP250 mutation is associated with CRD and hearing loss [2]; mutations in CNGB1 cause CRD and anosmia [3]; and CEP19 mutation leads to CRD with obesity and renal malformation[4]. Different from photoreceptor connecting cilia, sperm flagellum is motile cilia. Motile and immotile cilia defects usually do not occur at the same time. However, occasionally, CRD can also occur as a syndrome in combination of abnormalities of sperm flagella. As this syndrome leads to defects of two unrelated physiological functions, vision and reproduction, comprehensive analysis of its specific genetic etiology and molecular mechanism is still limited.

Centrosomal protein of 78 kDa (CEP78), protein encoded by the *CEP78* gene (MIM: 617110), is a centriolar protein composed of 722 amino acids and possesses two leucine-rich repeats and a coiled-coil domain[5]. CEP78 localizes to the distal region of mature centrioles and is exclusively expressed in ciliated organisms, supporting its crucial function in maintaining centrosome homeostasis[6, 7]. Mutations in the *CEP78* gene cause autosomal recessive CRD and hearing loss (CRDHL; MIM: 617236)[8–12]. Deregulation of Cep78 is also associated with colorectal cancer[13], prostate cancer[14], and asthenoteratozoospermia[12]. As a centrosomal protein, CEP78 functions downstream of CEP350 to control biogenesis of primary cilia [15], and regulates centrosome homeostasis through interactions with other centrosomal proteins, including CEP76, CP110 and EDD-DYRK2-DDB1^VprBP^[6]. Despite these findings, the biological function of Cep78, especially its function and mechanism in CRD with male infertility remains to be elucidated.

In this study, based on results of a male patient carrying *CEP78* mutation and *Cep78* gene knockout mice, we report CEP78 as a new causative gene for a distinct syndrome involving two phenotypes, CRD and male sterility[16]. We revealed that CEP78 deficiency caused dysfunction and irregular ciliary structure of photoreceptors. Deletion of Cep78 triggered male infertility, spermatogenesis defects, aberrant sperm flagella structures and manchette formation, and abnormal spermatid centriole length in male mice. The male patient carrying homozygous *CEP78* mutation leading to CEP78 protein loss also presented syndromic phenotypes including CRD, hearing loss, and multiple morphological abnormalities of the sperm flagella (MMAF). IFT20 and TTC21A are both IFT proteins have interaction with each other and vital for spermatogenesis and cilliogenesis[17]. CEP78 forms a trimer with IFT20 and TTC21A, which are vital for spermatogenesis and cilliogenesis[17–19], and regulated their interaction and stability. CEP78 recruits centrosome localization of IFT20. Defects of CEP78 trimer leaded to extended centriole and cilia shortening.

## Results

### Generation of *Cep78^-/-^* mice

To investigate the role of CEP78, we generated *Cep78* knockout mice using clustered regularly interspaced short palindromic repeats (CRISPR)/Cas 9 system. A sgRNA targeting exons 2 and 11 of the *Cep78* mice gene (**Supplementary Figure S1A**) was designed, generated, and injected into C57BL/6 mice zygote with the Cas 9 RNA. Genotyping confirmed deletion of exons 2 to 11 in heterozygotes and homozygotes (**Supplementary Figure S1B**). As revealed by immunoblotting, expression of Cep78 protein was completely absent in the neural retina and testes of *Cep78^-/-^* mice (**Supplementary Figures S1C-D**), suggesting successful generation of *Cep78* knockout mice.

### *Cep78^-/-^* mice show photoreceptor impairments

To reveal the retinal phenotypes associated with Cep78 deletion, we studied the functional and morphological changes in *Cep78^-/-^* mice retina. We initially used electroretinogram (ERG) to detect visual functions of *Cep78^-/-^* mice. ERG results of *Cep78^-/-^* mice and their heterozygous littermates at ages of 3-, 6-, 9-and 18-month were recorded. Scotopic a- and b-wave, together with photopic b-wave amplitudes were reduced in *Cep78^-/-^* mice when compared to age-matched *Cep78^+/-^* mice (**Figures 1A-E**). The reduction extended as age increased. According to our data, amplitudes of scotopic a-wave showed approximately 13.6%, 39.1%, 40.8%, and 64.8% decrease in *Cep78^-/-^* mice at 3-, 6-, 9- and 18-month respectively (**Figures 1A, C**), and amplitudes of scotopic b-wave were down-regulated by 19.9%, 30.5%, 35.6%, and 58.3% in *Cep78^-/-^* mice at 3-, 6-, 9- and 18-month respectively (**Figures 1A**, **D**). Similarly, amplitudes of photopic b-wave were decreased by 39.4%, 39.8%, 52.8%, and 69.0% in *Cep78^-/-^* mice at 3-, 6-, 9- and 18-month respectively (**Figures 1B, E**).

**Figure 1.**
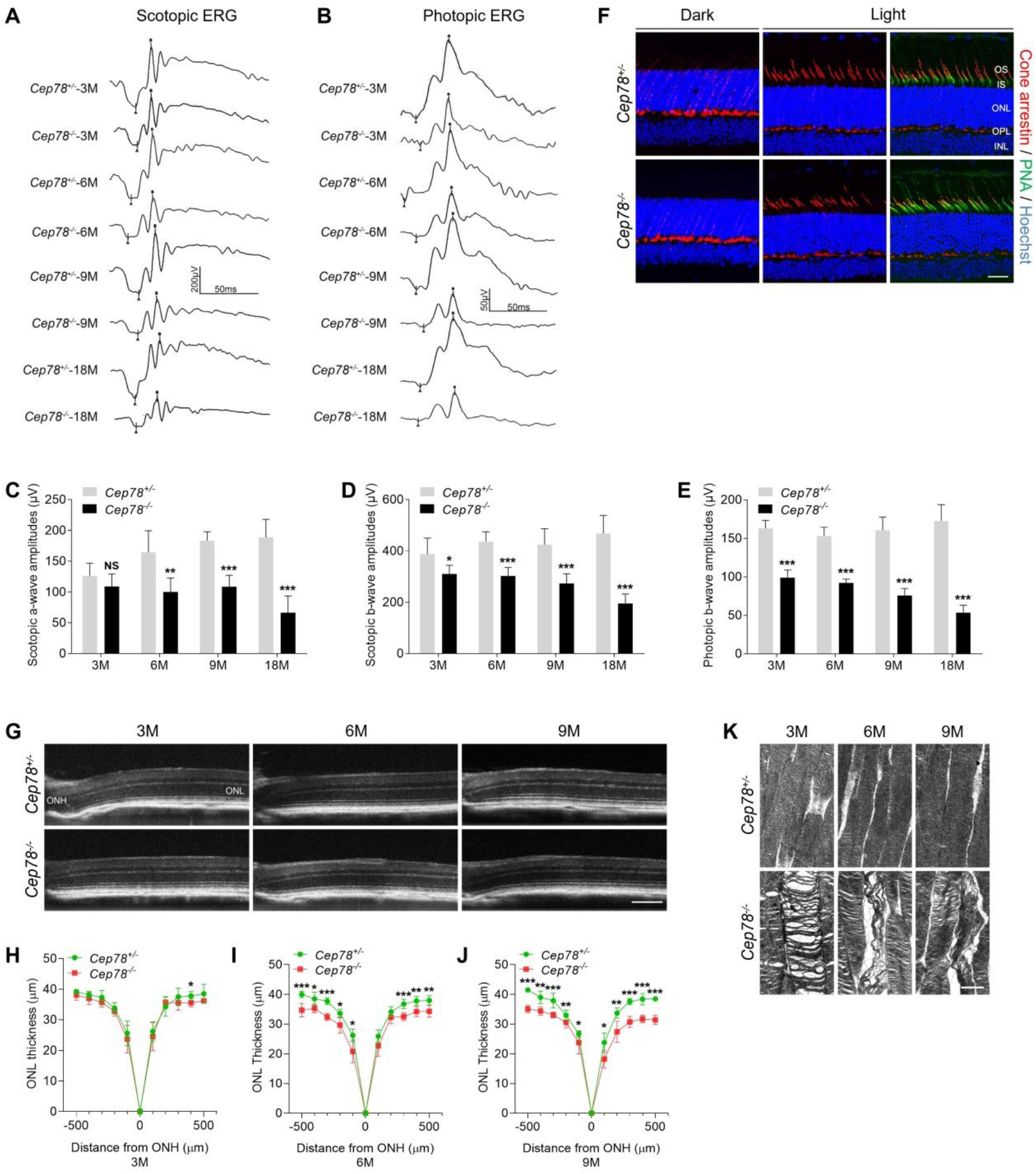
*Cep78* deletion leads to photoreceptor impairments in mice retina. (A-E) Scotopic and photopic ERG of *Cep78^-/-^* mice at the indicated ages. Representative images along with the quantification results were shown (n=6 for each sample, two-tailed Student’s t test). NS, not significant. *, *p* < 0.05, **, *p* < 0.01, ***, *p* < 0.001. (F) Immunofluorescence staining showing expression patterns of cone arrestin (CAR) and peanut agglutinin (PNA) lectin in dark- and light-adapted retinas of *Cep78^+/-^* and *Cep78^-/-^* mice at 3-month. Scale bar, 25 µm. (G-J) SD-OCT revealed thickness of outer nucleic layer (ONL) in retinas of *Cep78^+/-^* and *Cep78^-/-^* mice of indicated ages. Scale bar, 150 µm. Representative images along with the quantification results were shown (n=6 for each sample, two-tailed Student’s t test). *, *p* < 0.05, **, *p* < 0.01, ***, *p* < 0.001. (K) TEM revealed ultra-structures of photoreceptors in *Cep78^+/-^* and *Cep78^-/-^* mice at the indicated ages. Scale bar, 1 µm.

In normal retina, light exposure drives rapid movement of cone arrestin into outer segments (OS) of cone photoreceptors[20]. Herein, we tested whether Cep78 knockout would disrupt such translocation of cone arrestin in cone photoreceptors. Immunostaining patterns of cone arrestin in dark- and light-adapted *Cep78^+/-^* and *Cep78^-/-^* mice retinas were compared. In dark-adapted retina, cone arrestin was localized in the synaptic region, outer nuclear layer (ONL), and outer and inner segments. No difference was observed in distribution patterns of cone arrestin between *Cep78^+/-^* and *Cep78^-/-^* mice (**Figure 1F**). Light exposure of 2 hours triggered a shift of cone arrestin from the inner cellular compartments to the OS in both *Cep78^+/-^* and *Cep78^-/-^* mice. However, light-induced translocation of cone arrestin in *Cep78^-/-^* mice retina was slower and more limited than in *Cep78^+/-^* mice (**Figure 1F**). Thus, above data suggested that regular functions of retina were disturbed in *Cep78^-/-^* mice.

We next investigated morphological changes in retina of *Cep78^-/-^* mice. Spectral domain-optical coherence tomography system (SD-OCT) was applied to visualize all layers of mice retina. SD-OCT showed that thicknesses of ONL were attenuated in *Cep78^-/-^* mice when compared to *Cep78^+/-^* mice at ages of 3-, 9-, and 12-month (**Figures 1G-J**). The attenuation became more evident along with increasing of age (**Figures 1G-J**). We also utilized transmission electron microscopy (TEM) to observe the ultra-structure of photoreceptors. As evidenced by TEM, photoreceptor OS are regularly lined and shaped in *Cep78^+/-^* mice at 3-, 6-, and 9-month age. However, disorganized, sparse, and caduceus photoreceptor OS disks, and widen OS bases were observed in age-matched *Cep78^-/-^* mice (**Figure 1K**). Collectively, our results indicated that both functions and morphologies of mice photoreceptor were disturbed upon Cep78 deletion.

### *Cep78^-/-^* mice exhibit disturbed ciliary structures in photoreceptors

Since CEP78 is a ciliary protein and *CEP78* mutation was found associated with primary-cilia defects[9], we then aimed to analyze whether *CEP78^-/-^* mice exhibit abnormal ciliary structure using immunofluorescence staining and TEM. Ciliary axonemes and basal bodies were labeled with antibody against Ac-α-tubulin in immunofluorescence staining. Our data indicated that cilia were shortened in *Cep78^-/-^* mice at 3-, 6-, and 9-month age compared to age-matched *Cep78^+/-^* mice (**Figure 2A**). We further used TEM to visualize longitudinal sections of the ciliary region of photoreceptors. Based on our findings, photoreceptors of *Cep78^-/-^* mice had shortened connecting cilia with their microtubules spread in their upper part (**Figures 2B-F**). Structures of basal bodies, adjacent daughter centrioles, or other organelles of inner segment were not altered in *Cep78^-/-^* mice (**Figures 2D-F**). Thus, our data suggested that Cep78 deficiency interrupts cilia elongation and disturbed ciliary structures in photoreceptors.

**Figure 2.**
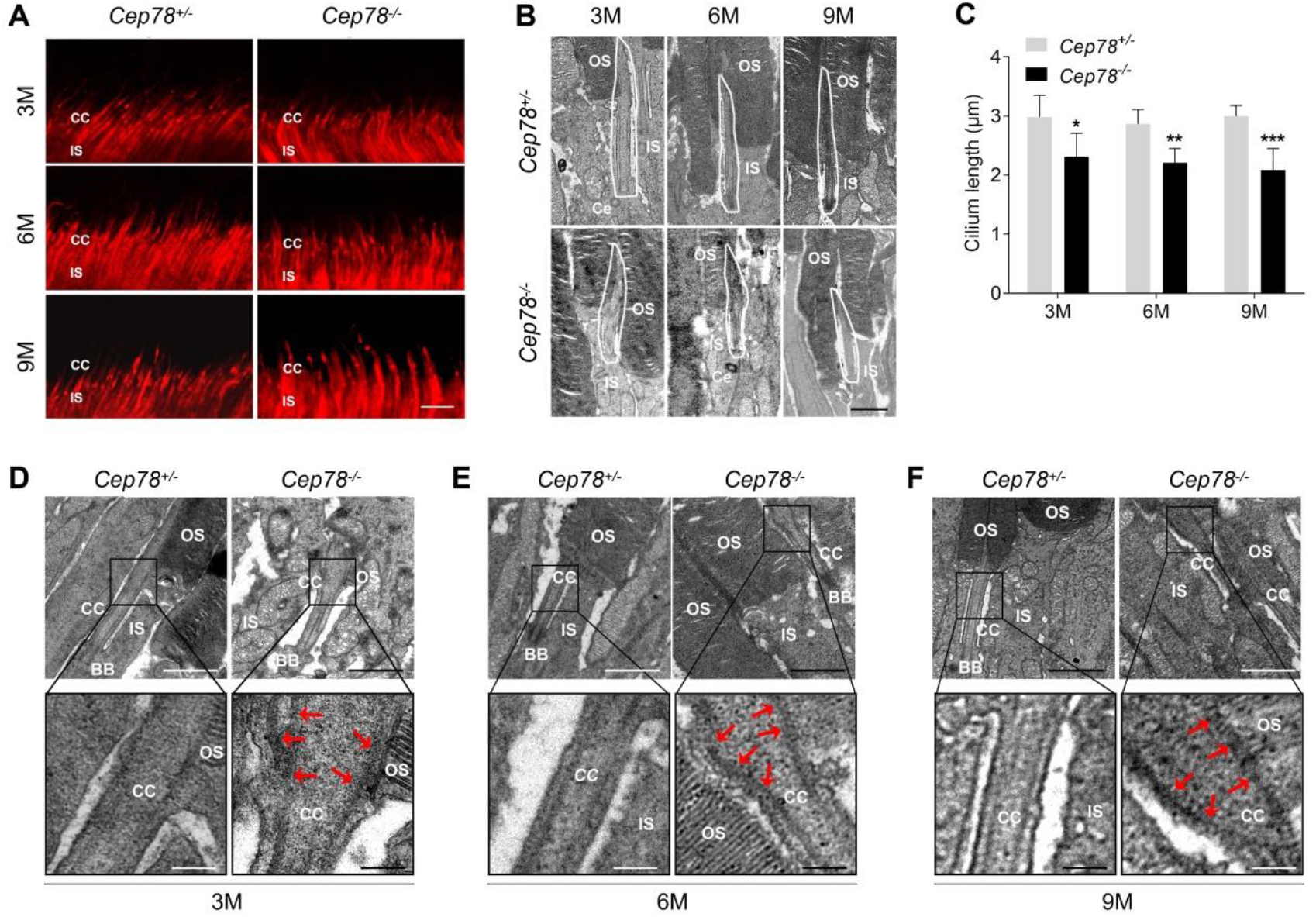
*Cep78^-/-^* mice exhibit disturbed ciliary structure in photoreceptors. (A) Retinal cryosections from *Cep78^+/-^* and *Cep78^-/-^* mice at indicated ages stained with Ac-α-tubulin (red) to visualize ciliary axonemes and basal bodies. Scale bar, 5 µm. CC, connecting cilium; IS, inner segment. (B-C) TEM was applied to observe longitudinal sections of the ciliary region of photoreceptors in *Cep78^+/-^* and *Cep78^-/-^* mice at indicated ages. Scale bar, 1 µm. OS, outer segment; IS, inner segment; Ce, Centriole. n=6 for each sample, *, *p* < 0.05, **, *p* < 0.01, ***, *p* < 0.001. (D-F) TEM was used to observe the ultrastructure of connecting cilium in photoreceptors of *Cep78^+/-^* and *Cep78^-/-^* mice at indicated ages. Scale bar, upper 1 µm, below 200 nm. OS, outer segment; CC, connecting cilium; IS, inner segment; BB, basal body.

### Cep78 deletion leads to male infertility and MMAF in mice

Unexpectedly, we found that *Cep78^-/-^* male mice were infertile during the breeding of *Cep78^-/-^* mice. We thus explored the reproductive functions of *Cep78^-/-^* male mice. No difference was revealed between the testis weight, body weight, and testis/body weight of *Cep78^+/-^* and *Cep78^-/-^* male mice (**Supplementary Figures S2A-C**). Mating test with *Cep78^+/-^* female mice for 3 months showed that *Cep78^-/-^* male mice were infertile (**Figure 3A**). To further explore the underlying causes for male infertility of *Cep78^-/-^* mice, we assessed the concentration, motility and progressive motility of sperm isolated from cauda epididymis using computer-assisted sperm analysis (CASA), and all three parameters were decreased upon *Cep78* deletion (**Supplementary Table S1** and **Supplementary Movies 1-2**). Consistently, hematoxylin-eosin (H&E) staining also showed less spermatozoa in both caput and cauda epididymis of *Cep78^-/-^* male mice compared with age matched *Cep78^+/-^* male mice (**Figure 3B**).

**Figure 3.**
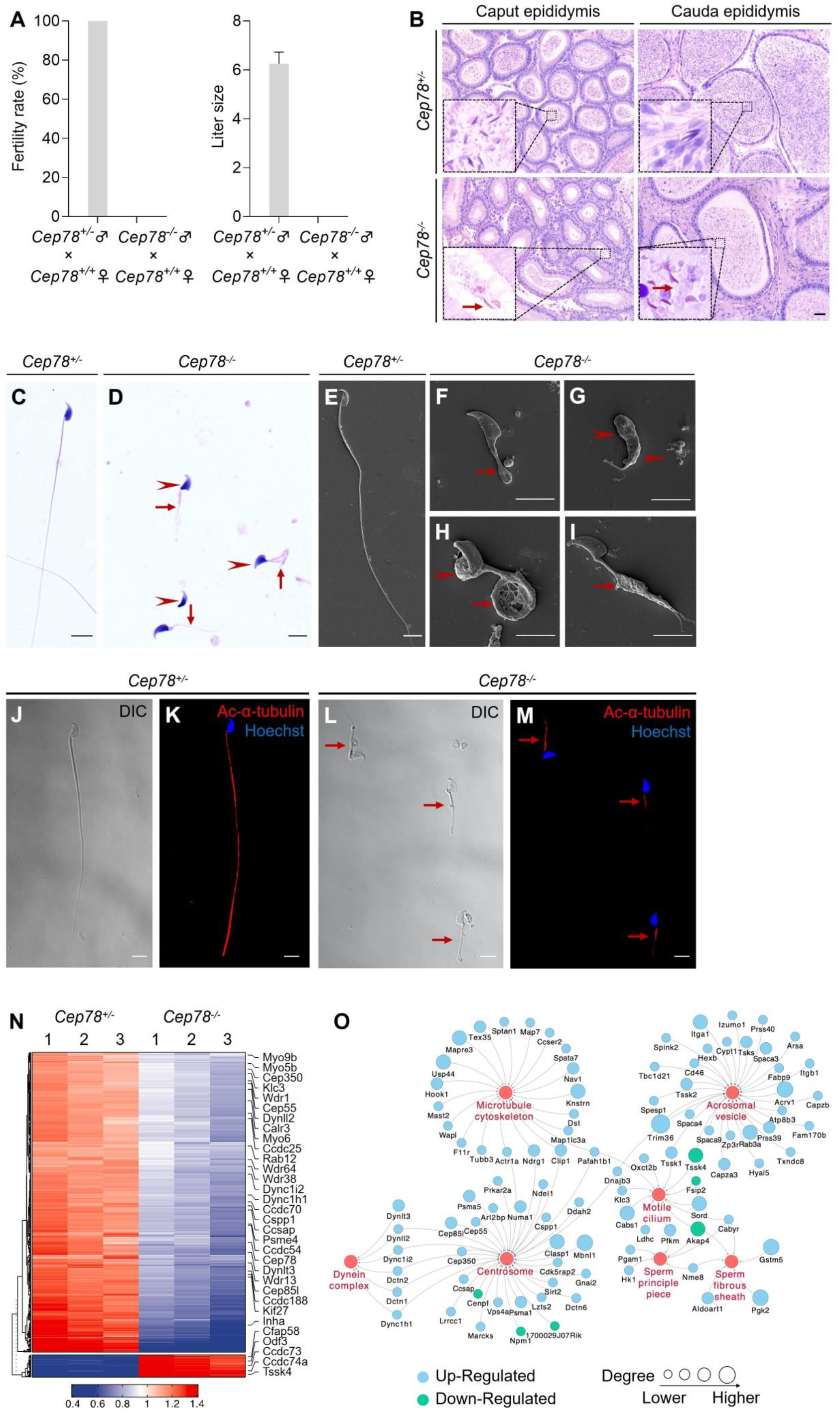
*Cep78^-/-^* mice present male infertility and morphological abnormalities of sperm. (A) Fertility rates (left) and litter sizes (right) of *Cep78^+/-^* and *Cep78^-/-^* male mice crossed with *Cep78^+/+^* female mice, n=4 for *Cep78^+/-^* male mice crossed with *Cep78^+/+^* female mice, n=5 for *Cep78^+/-^* male mice crossed with *Cep78^+/+^* female mice. (B) H&E staining was utilized to observe the structures of caput and cauda epididymis in *Cep78^+/-^* and *Cep78^+/-^* mice. Scale bar, 100 µm. (C-D) Spermatozoa from *Cep78^+/-^* (C) and *Cep78^-/-^* (D) mice were observed with H&E staining. Abnormal heads and short, missing or coiled flagella were indicated by arrow heads and arrows respectively. Scale bar, 5 µm. (E-I) Spermatozoa of *Cep78^+/-^* (E) and *Cep78^-/-^* (F-I) mice were observed by SEM. Abnormal heads were represented by arrow heads. Absent, short, coiled or multi flagella were indicated by arrows. (J-M) Spermatozoa of *Cep78^+/-^* and *Cep78^-/-^* mice were pictured using differential interference contrast (J, L), and were stained with Ac-α-tubulin (red) and Hoechst (blue) to visualize axonemes and heads, respectively (K, M). (N) Heatmap showing differentially expressed proteins between *Cep78^+/-^* and *Cep78^−/−^* mice elongating spermatids. (O) Network analysis of GO cellular components terms enriched in differential expressed proteins between *Cep78^+/-^* and *Cep78^−/−^* mice elongating spermatids. Scale bar, 5μm.

Multiple abnormalities of sperm head and flagella in *Cep78^-/-^* male mice were initially detected by light microscopy (**Figures 3C-D**) and scanning-electron microscopy (SEM; **Figures 3E-I**), which identified severely distorted sperm heads and flagella (**Figures 3D, F-I**), including absent (**Figures 3D, F**), short (**Figures 3D, G**), coiled (**Figures 3D, H**), and multi-flagella (**Figure 3I**). Spermatozoa with abnormal heads, necks and flagella accounted for 92.10%, 82.61% and 95.67% of total spermatozoa respectively in *Cep78^-/-^* male mice (**Supplementary Table S1**). Immunofluorescence staining was also used to study sperm morphology, results of which were consistent with those of light microscopy and SEM. Sperm axoneme was stained using antibody against Ac-α-tubulin, and short flagella were frequently observed (**Figures 3J-M**).

We next applied quantitative mass spectrometry (MS) on elongating spermatids lysates of *Cep78^+/-^* and *Cep78^-/-^* mice to reveal effects of Cep78 insufficiency on testicular protein expressions and pathway regulations. We set the threshold as a fold-change greater than 1.5 and p value less than 0.05, and identified a total of 806 down-regulated proteins and 80 up-regulated proteins upon Cep78 depletion, which were visualized using hierarchical clustering analyses (**Figure 3N** and **Supplementary Table S2**). Consistent to above findings in morphological anomalies in sperm, significantly enriched gene ontology (GO) terms in cellular components included sperm principal piece, sperm fibrous sheath, acrosomal vesicle, motile cilium, centrosome, microtubule skeleton and dynein complex (**Figure 3O** and **Supplementary Table S3**). Taken together, Cep78 deletion leaded to decreased count, motility and progressive motility of sperm, morphological abnormalities of sperm head and flagella in mice, as well as dysregulated sperm proteins thus accounting for male infertility.

### Defective microtubule arrangements and elongated centrioles in sperm flagella of *Cep78^-/-^* mice

To determine which spermatogenesis stage that Cep78 affected on, we compared histology of testicular tissue sections between *Cep78^-/-^* and *Cep78^+/-^* male mice using periodic acid-Schiff (PAS) staining. Light microscopy showed an apparent scarcity of sperm flagella visible in the lumens of *Cep78^-/-^* testes (**Figures 4A-B**), with flagellar formation defects observed from stages I-III till stages VII-VIII during spermatogenesis (**Figures 4B**), suggesting that Cep78 was involved at the stage of sperm flagellar formation. Very few spermatids reached maturity successfully and passed into epididymis (**Supplementary Table S1**).

**Figure 4.**
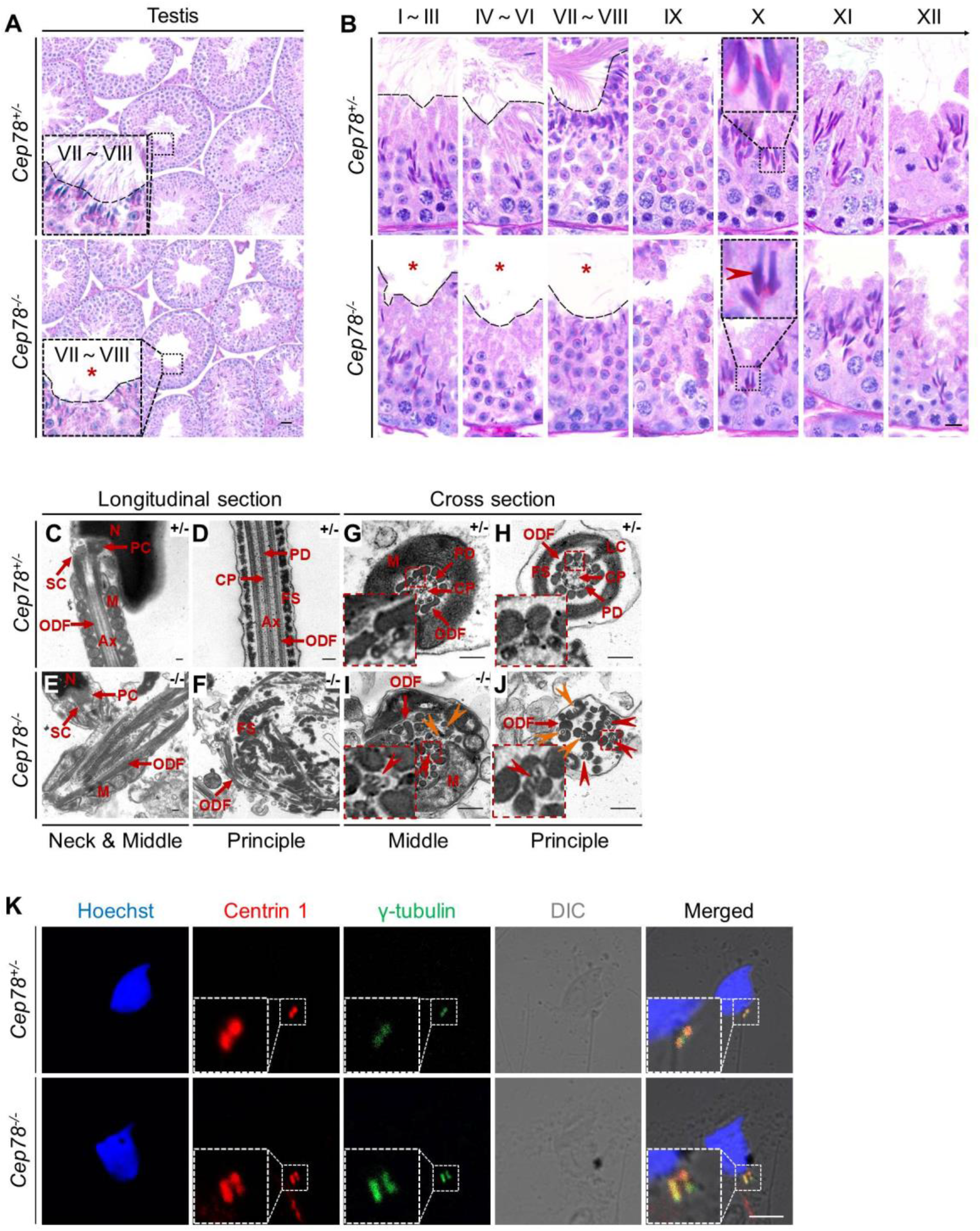
Defective microtubule arrangements and elongated centrioles in sperm flagella of *Cep78^-/-^* mice. (A) Paraffin sections of testicular tissues from *Cep78^+/-^* and *Cep78^-/-^* male mice were stained with PAS. Defects of sperm tails were indicated by asterisk. Scale bar, 50μm. (B) Twelve stages of spermatogenesis in *Cep78^+/-^* and *Cep78^-/-^* male mice were presented by PAS-stained paraffin sections of seminiferous tubules. Lack of sperm tails and abnormal nuclei shape of elongated spermatids were indicated by asterisk and arrow head, respectively. Schematic diagrams were attached. Scale bar, 10μm. (C-J) TEM was applied to visualize ultra-structures of *Cep78^+/-^* and *Cep78^-/-^* spermatozoa in longitudinal sections of neck & middle pieces (C and E) and principal pieces (D and F), and in cross sections of middle pieces (G and I) and principal pieces (H and J). Triplet and singlet microtubules were indicated by red and orange arrow heads (I-J), respectively. Scale bar, 200 nm. (K) *Cep78^+/-^* and *Cep78^-/-^* mice spermatozoa were stained with centrin 1 and γ-tubulin to reveal the structure of centrosome. Scale bar, 5 µm.

To further carefully characterize *Cep78^-/-^* mice phenotype, we analyzed the ultra-structure of mouse testes using TEM. Based on our findings, *Cep78^+/-^* sperm showed well aligned arrangements of connecting segments, axoneme microtubules, mitochondria, outer dense fibers (ODF), and fiber sheaths in longitudinal section of neck, middle, and principal pieces (**Figures 4C-D**). However, disordered mitochondria and a jumble of unassembled flagellar components were presented in *Cep78^-/-^* sperm (**Figures 4E-F**). In cross sections of middle and principal regions, *Cep78^+/-^* sperm exhibited well-aligned ODF and typical “9+2” arrangement of microtubules, which were typified by 9 peripheral microtubule doublets surrounding a central pair of singlet microtubules (**Figures 4G-H**). In contrast, *Cep78^-/-^* sperm showed complete derangement of ODF and scattered “9+2” microtubules (**Figures 4I-J**). In normal sperm flagella, distal centriole develops an atypical centriole structure, whose microtubules are doublets instead of triplets. Triplet microtubules can only be observed at proximal centrioles[21]. In *Cep78*^-/-^ sperm, triplet microtubules, which were supposed to exist only in proximal centrioles, were found at both middle and principal pieces of sperm flagella (**Figures 4I-J**), indicating abnormal elongation of centrioles during sperm flagella formation.

Tubules in axoneme are formed from centrosome. The various types of aberrant sperm flagella observed in mice with CEP78 defects, together with the triplet microtubules found in principal and middle pieces of sperm flagella upon CEP78 deletion, suggested centriole anomalies as potential causes for deregulation of sperm development in mouse. Previous *in vitro* studies showed that interference of Cep78 leaded to elongation of centriole[6]. To further elucidate whether Cep78 regulates centriole length in spermiogenesis, we stained centrioles and peri-centriole materials with anti-centrin 1 and γ-tubulin, respectively. Two-dot staining pattern of disengaged centrioles was observed in *Cep78^+/-^* spermatids (**Figure 4K**), while depletion of Cep78 promoted centriole elongation in spermatids (**Figure 4K**), supporting the role of Cep78 in regulating centriole structures and functions. Collectively, our data revealed defective microtubule arrangements and elongated centrioles in sperm flagella of *Cep78^+/-^* mice.

### *Cep78^-/-^* testes present abnormalities in spermatid head formation during spermiogenesis

In addition to axonemal defects in sperm flagella, *Cep78^-/-^* testes also exhibit deformation of spermatid nuclei, which was presented by abnormal club-shaped nuclear morphology of elongated spermatids since step 10, in contrast to the normal hook-shaped head in *Cep78^+/-^* elongated spermatids (**Figures 4B, 5A**). Abnormal elongation of manchette structures in testicular spermatids of *Cep78^-/-^* mice during spermiogenesis was also shown by immunofluorescence staining (**Figure 5B**), which supported above findings by PAS staining.

**Figure 5.**
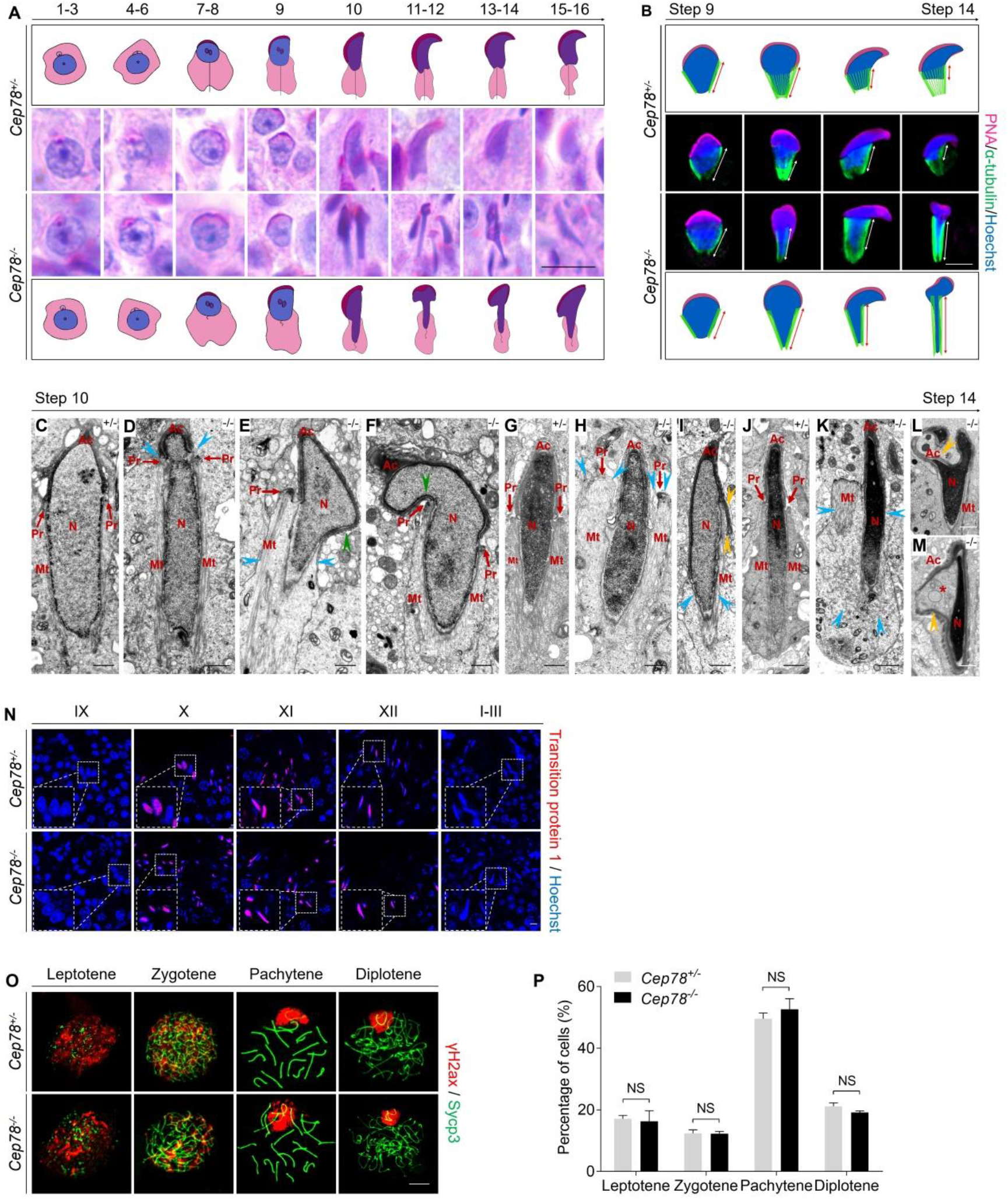
*Cep78^-/-^* testes present abnormalities in spermatid head formation during spermiogenesis. (A) Sixteen steps of spermiogenesis in *Cep78^+/-^* and *Cep78^-/-^* male mice were presented by PAS-stained paraffin sections of seminiferous tubules. Lack of sperm tails and abnormal nuclei shape of elongated spermatids were indicated by asterisk and arrow head, respectively. Schematic diagrams were attached. Scale bar, 10μm. (B) Elongated spermatids from stage 9-14 from *Cep78^+/-^* and *Cep78^-/-^* mice were stained with PNA (red) and α-tubulin (green) to see elongation of manchette structures. Nuclei were counterstained with Hoechst (blue). Manchette structures were recognized by double-head arrows. Scale bar, 5μm. Schematic diagrams were attached. (C-M) TEM was used to visualize ultra-structures of *Cep78^+/-^* and *Cep78^-/-^* spermiogenic spermatids from step 10-14. Scale bar, 500 nm. N, nucleus; M, mitochondria; Ax, axoneme; SC, segmented column of the connecting piece; PC, proximal centriole; ODF, outer dense fiber; PD, peripheral microtubule doublets; CP, central pair of microtubules; FS, fibrous sheath; LC, longitudinal column; Ac, acrosome; Pr, perinuclear ring; Mt, manchette. Abnormal structures of manchette were indicated by celeste arrow heads (D, E, H, I, K). Green arrow heads pointed at abnormal bend of spermatid heads (E, F). Yellow arrow heads represented abnormal acrosomes (I, L, M). Asterisk indicated expanded perinuclear space (M). (N) Paraffin sections of testicular seminiferous tubules from stage IX of spermatogenesis to stage I in *Cep78^+/-^* and *Cep78^-/-^* mice were stained with transition protein 1 (red) and Hoechst (blue) to observe nuclear condensation. Scale bar, 5μm. (O-P) Immunofluorescence staining of γH2ax (red) and Sycp3 (green) in chromosome spreads of spermatocytes from the testes of *Cep78^+/-^* and *Cep78^-/-^* mice. Scale bar, 5μm. Representative image along with the quantification results was shown (n=219 for Cep78^+/-^ spermatocytes, n=208 for Cep78^-/-^ spermatocytes, two-tailed Student’s t test, NS: not significant).

Further analyses with TEM observed various ultra-structural defects in sperm head formation of *Cep78*^-/-^ spermatids. As indicated by TEM, *Cep78*^+/-^ spermatids underwent a series of dramatic changes and presented regular reshaping of sperm head, including nuclear condensing, manchette formation, and acrosomal biogenesis (**Figures 5C**, **G, J**). TEM suggested that nuclear condensing was normal in *Cep78*^-/-^ spermatids (**Figures 5D-F, H-I, K-M**). Similar expressional intensity and localization of transition protein 1 from stage IX of spermatogenesis to stage I-III in *Cep78*^+/-^ and *Cep78*^-/-^ mice were revealed by immunofluorescence staining, confirming the TEM data and suggesting that nuclear condensation was not affected upon Cep78 deletion (**Figure 5N**). We also performed spermatocyte spreading of *Cep78*^+/-^ and *Cep78*^-/-^ mice testicular tissues to detect ratios of spermatocytes at leptotene, zygotene, pachytene, and diplotene stages. Based on our results, Cep78 loss showed no effects on spermatocyte development at all four stages (**Figures 5O-P**), implying that Cep78 deletion did not affect the prophase of meiosis I.

The manchette structure, consisted of a series of parallel microtubule bundles that extended from perinuclear rings of nucleus to distal cytoplasm and are closely proximity to or parallel to nuclear membranes, regulates sperm head formation during spermiogenesis[22]. We herein revealed various abnormalities in manchette formation and acrosomal biogenesis in *Cep78*^-/-^ spermatids, including an abnormal nuclear constriction at the site of perinuclear rings (**Figure 5D**), ectopic and asymmetric perinuclear rings together with disordered manchette microtubules (**Figures 5E-F, H-I, K**), abnormal acrosome and nuclei invagination (**Figure 5L**), and detached acrosome from nuclear membrane (**Figure 5M**). The defective spermatid manchette formation in *Cep78^-/-^* mice led to peculiar shaping of nucleus and abnormal sperm head formation. Therefore, the above findings indicated that absence of Cep78 generates defective spermatid head formation.

### *CEP78* mutation causes male infertility and MMAF in human patient with CRD

We have previously linked the *CEP78* c.1629-2A>G mutation with autosomal recessive CRDHL, and revealed a 10 bp deletion of *CEP78* exon 14 in mRNA extracted from white blood cells of patient carrying *CEP78* c.1629-2A>G mutation[8]. We further explored whether this mutation will disturb CEP78 protein expression using immunoblotting. Blood sample was collected from this patient and an unaffected control for protein extraction. Our results revealed absence of CEP78 protein in white blood cells of patient carrying *CEP78* c.1629-2A>G mutation (**Supplementary Figure S3**), suggesting that this mutation leads to loss of CEP78 protein.

Since Cep78 deprivation led to male infertility and sperm flagellar defects in mice, and *CEP78* c.1629-2A>G mutation caused degradation of CEP78 protein in human, we thus investigated reproductive phenotype of the previously reported male patient, who carried homozygous *CEP78* c.1629-2A>G mutation and was diagnosed with CRD and hearing loss[23]. The patient is infertile. His semen volume was 2.6 mL, which was among the normal range (**Supplementary Table S1**). However, CASA revealed obvious reductions in sperm concentration, motility, and progressive motility in the *CEP78*-mutated man according to standard of the World Health Organization (WHO) guidelines (**Supplementary Table S1**). In addition, impaired sperm movement of the patient is presented in **Supplementary Movies 3**.

Patient’s semen sample was further submitted for morphological analyses using light microscopy and SEM, both of which showed typical MMAF phenotypes, such as abnormal heads (95.42%), necks (85.62%) and flagella (97.39%) (**Figures 6A-J** and **Supplementary Table S1**). A great diversity of abnormal flagella was observed including short flagella (**Figures 6B, G**), coiled flagella (**Figures 6C, I**), absent flagella (**Figures 6D, H**), and multi flagella (**Figures 6E, J**). The patient’s sperm also had a series of sperm head abnormalities, such as large acrosome area (**Figure 6B**), pear shaped nucleus (**Figure 6C**), conical nucleus (**Figure 6D**), and excessive acrosome vacuoles (**Figure 6E**). TEM was further applied to visualize the ultra-structures of spermatozoa in the male patient carrying *CEP78* c.1629-2A>G mutation. Based on our data, multiple ultra-structural abnormalities were observed in the spermatozoa of case as compared to those of age matched healthy male individual (**Figures 6K-R**). For example, longitudinal sections showed disordered arrangements of mitochondrial sheaths, severe axonemal disorganization, and fibrous sheath hyperplasia (**Figures 6L, N**). Instead of the typical “9+2” microtubule structure in exoneme of normal sperm flagella (**Figures 6O, Q**), absent or reduced central-pair microtubules and disarranged peripheral microtubule doublets were frequently observed in cross sections (**Figures 6P, R**). Similar to the *Cep78*^-/-^ mice data, triplet microtubules unique to proximal centrioles in sperm were also observed in both principal and middle pieces of patient’s sperm flagella (**Figures 6P, R**). Taken together, this patient carrying *CEP78* c.1629-2A>G mutation presented typical MMAF phenotypes.

**Figure 6.**
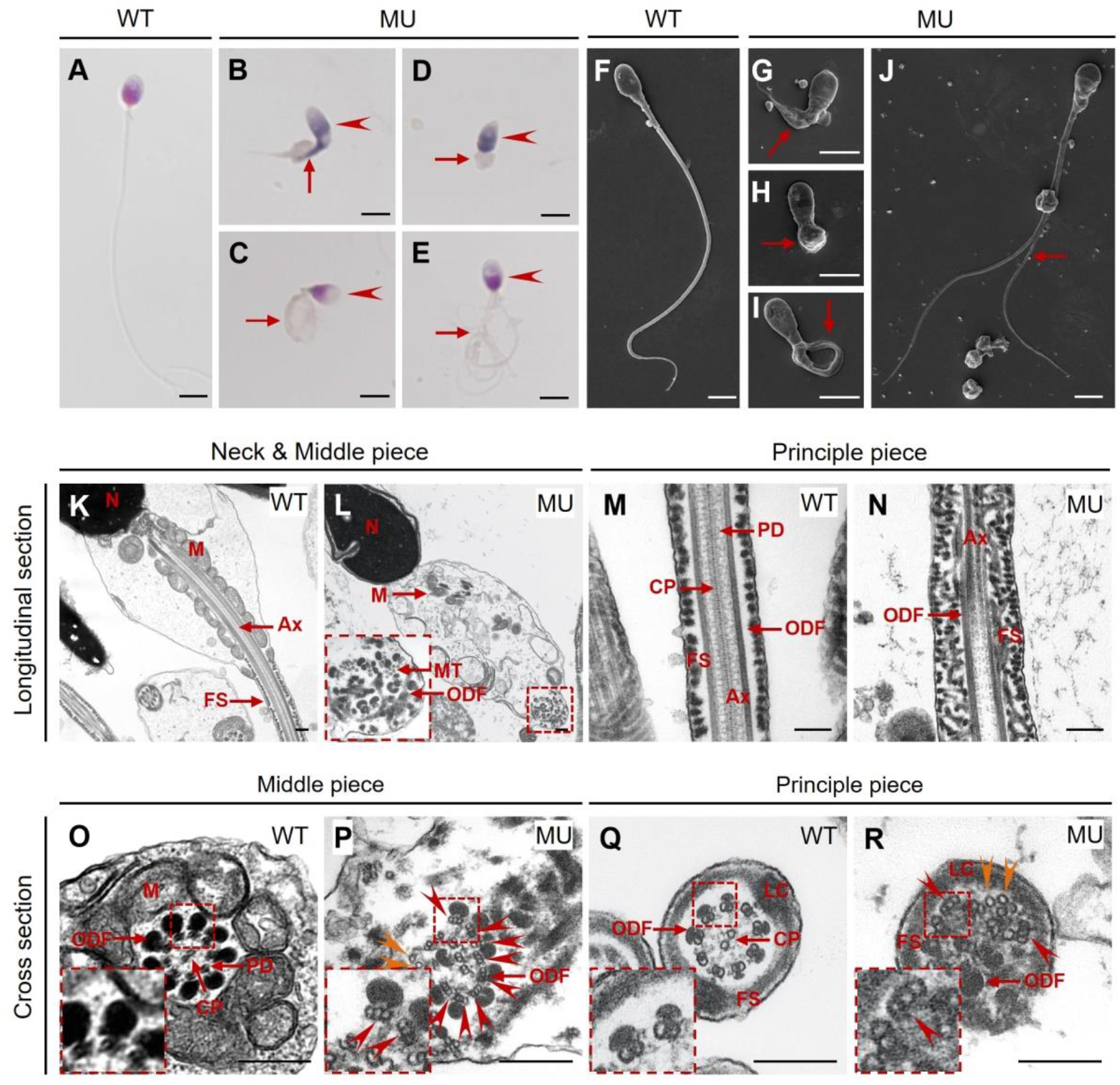
*CEP78* mutation causes MMAF phenotypes in human. (A-E) HE staining was utilized to see structures of spermatozoa from a healthy control (A) and a patient carrying homozygous *CEP78* c.1629-2A>G mutation (B-E). Scale bar, 5μm. Arrow heads represented abnormal sperm heads. Short, coiled, absent or multi flagella were indicated by arrows. (F-J) Ultra-structures of spermatozoa from the healthy control (F) and the patient with *CEP78* mutation (G-J) were observed using SEM. Short, coiled, absent or multi flagella, and cytoplasm remain were indicated by arrows. Scale bar, 5μm. (K-R) TEM was applied to visualize ultra-structures of spermatozoa from the healthy control and the patient in longitudinal sections of neck & middle pieces (K-L) and principal pieces (M-N), and in cross sections of middle pieces (O-P) and principal pieces (Q-R). Triplet and singlet microtubules were indicated by red and orange arrow heads (P, R), respectively. Scale bar, 200 nm (K-R). N, nucleus; M, mitochondria; MT, microtubules; Ax, axoneme; FS, fibrous sheath; ODF, outer dense fiber; PD, peripheral microtubule doublets; CP, central pair of microtubules; LC, longitudinal column.

### CEP78 forms a trimer with IFT20 and TTC21A, and is essential for their interaction and stability to regulate centriole and cilia lengths

To reveal the interaction between CEP78 and other proteins during spermiogenesis, we performed anti-Cep78 immunoprecipitation (IP) coupled with quantitative MS (IP-MS) on testicular lysates of *Cep78^+/-^* and *Cep78^-/-^* mice. A total of 16 ciliary proteins were initially screened out by proteomic analyses (**Supplementary Table S4**), among which Ift20 and Ttc21A were essential for sperm flagella assembly and male fertility. These two proteins bound together as a dimer and loss of IFT20 or TTC21A was found to cause male infertility and MMAF phenotypes[17, 19], we thus hypothesized that CEP78 regulates human and mice phenotypes by interacting with IFT20 and TTC21A as a tri-mer. To explore this hypothesis, we first tested whether the three proteins directly bind with each other using co-immunoprecipitation (co-IP) and immunoblotting analyses in HEK293T cells overexpressing tagged proteins. Direct interactions between CEP78 and IFT20 (**Figure 7A**), CEP78 and TTC21A (**Figure 7B**), and IFT20 and TTC21A (**Figure 7C**) were identified, supporting our hypothesis.

**Figure 7.**
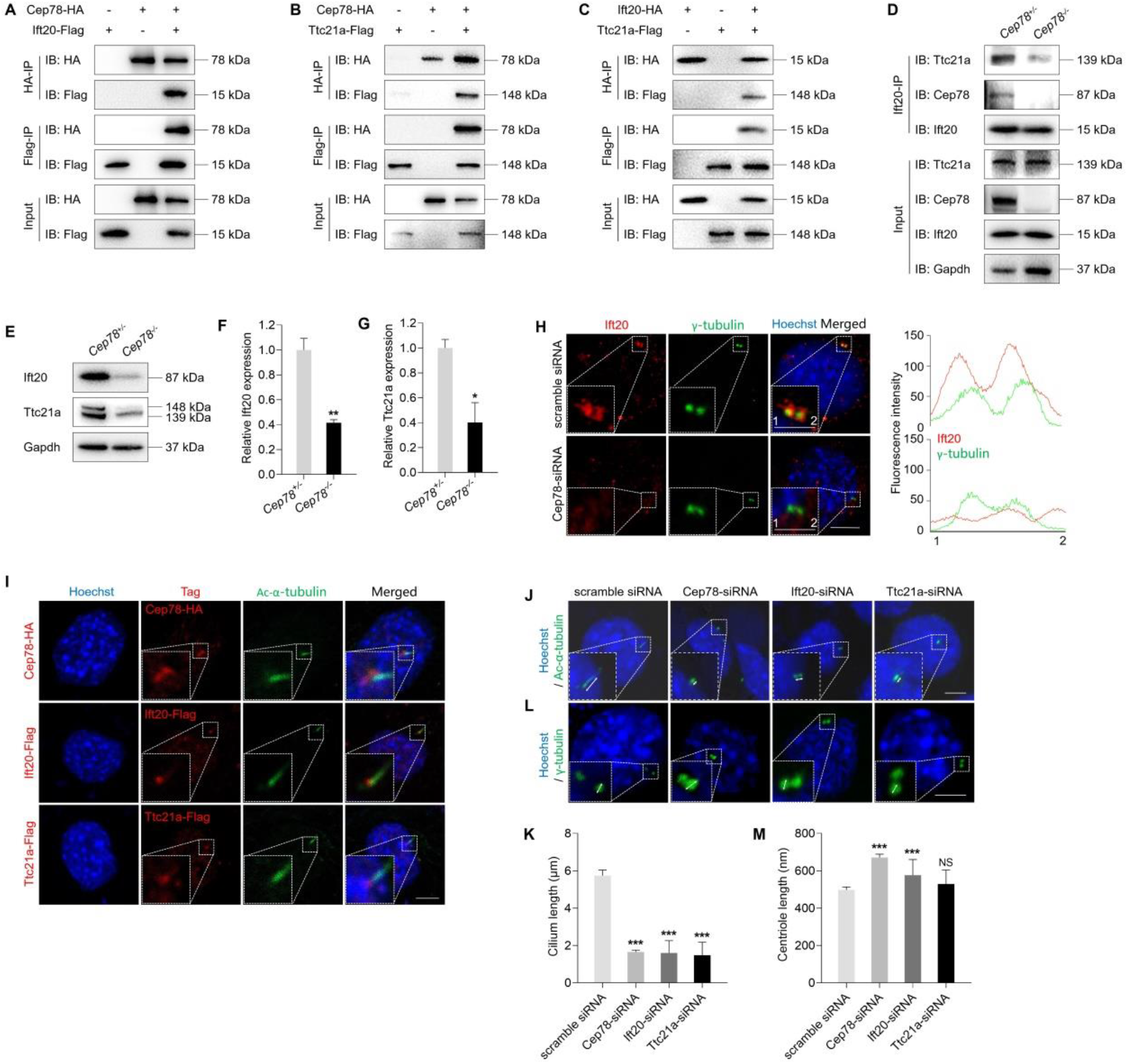
CEP78 forms a trimer with IFT20 and TTC21A to regulate axonemal and centrosomal elongations. (A-C) HEK293T cells were co-transfected with plasmids expressing Cep78-HA and Ift20-Flag (A), Cep78-HA and Ttc21a-Flag (B), or Ift20-HA and Ttc21a-Flag (C). Cellular lysates were immunoprecipitated with an anti-HA or anti-Flag antibody in 1% SDS and then immunoblotted (IB) with anti-HA or anti-Flag antibodies. (D) Lysates of testicular tissues from *Cep78^+/-^* and *Cep78^-/-^* male mice were immunoprecipitated with anti-Ift20 antibody in 1% SDS and then immunoblotted with anti-Ttc21a, anti-Cep78 or anti-Ift20 antibodies. (E-G) Immunoblotting showed Ift20 and Ttc21a expressions in lysates of testes from *Cep78^+/-^* and *Cep78^-/-^* mice (E). Representative images along with the quantification results were shown (F and G; 3 biological replications for *Cep78^+/-^* and *Cep78^-/-^* testes, two-tailed Student’s t test). (H) 3T3 cells transfected with scramble siRNA or Cep78-siRNA were stained with Ift20 and γ-tubulin to reveal the intensities and locations of Ift20 and centrioles. Scale bar, 5 µm. A representative image of along with the quantification result was shown. (I) HA-tag, Flag-tag and cilia in 3T3 cells overexpressing Cep78-HA, Ift20-Flag or Ttc21a-Flag were visualized using antibodies against HA, Flag and Ac-α-tubulin, respectively. Nuclei were counterstained with Hoechst (blue). (J-M) Cilia (J) and centrioles (L) in 3T3 cells transfected with scramble siRNA, Cep78-siRNA, Ift20-siRNA or Ttc21a-siRNA were stained with Ac-α-tubulin and γ-tubulin respectively. Nuclei were counterstained with Hoechst (blue). Cilium and centriole structures were recognized by double-head arrows. Scale bar, 5μm. Representative images along with the quantification results were shown (K and M; Comparisons in cilia and centriole size between the scramble siRNA, Cep78 siRNA, Ift20 siRNA and Ttc21a siRNA were addressed by accumulated data from 3 independent experiments, for cilia length: n=35 for WT, n=34 for Cep78 siRNA,n=38 for Ift20 siRNA,n=40 for Ttc21a siRNA;for centriole length: n=21 for WT, n=25 for Cep78 siRNA,n=44 for Ift20 siRNA,n=55 for Ttc21a siRNA; one-way ANOVA, with Dunnet’s multiple comparison test amongst all groups). *, *p* < 0.05; **, *p* < 0.01; ***, *p* < 0.001; NS, not significant.

We next detected whether Cep78 depletion would cause instability of the protein complex and disturbs its functions. As revealed by co-IP and immunoblotting results, the direct binding between Ift20 and Ttc21a was severely disrupted upon Cep78 knocking out (**Figure 7D**), indicating that Cep78 is essential for maintaining stability of the tri-mer. We also found that expressions of both Ift20 and Ttc21a were decreased in testes of *Cep78*^-/-^ mice compared to *Cep78*^+/-^ mice (**Figures 7E, F, G**), suggesting that disruption of the trimer potentially affected stability of all components. Decreased Ift20 expression was also detected by immunofluorescence staining in 3T3 cells transfected with Cep78-siRNA compared to cells transfected with scramble siRNA (**Figure 7H**). Quantitative real-time PCR (Q-PCR) identified that expressions of Cep78 was remarkably knocked down to 15% by Cep78-siRNA in 3T3 cells (**Supplementary Figure S4A**). In addition, the co-localization between Ift20 and centrioles in 3T3 cells transfected with scramble siRNA was altered upon Cep78 knocking down (**Figure 7H**), supporting that Cep78 insufficiency might disturb the regular functions of Ift20.

We then aimed to determine the function of this tri-mer. We found that all three components of the trimer located in the base of the cilium of 3T3 cells (**Figure 7I**), indicating its role in regulating ciliogenesis. Ciliary axonemes and centrioles were labeled with antibodies to Ac-α-tubulin and γ-tubulin respectively. We found that cilium length was reduced in 3T3 cells transfected with Cep78-siRNA, Ift20-siRNA or Ttc21a-siRNA compared to cells transfected with scramble siRNA (**Figures 7J-K**). Efficiencies for Ift20 and Ttc21a knocking down using Ift20-siRNA and Ttc21a-siRNA in 3T3 cells were confirmed by Q-PCR (**Supplementary Figures S4B-C**). However, centrioles were elongated in 3T3 cells with Cep78 or Ift20 knocking down, but not in cells with insufficient Ttc21a (**Figures 7L-M**). Thus, our data suggested that insufficiency of any component in the trimer causes cilia shortening and centriole elongation, which was consistent with phenotypes we observed in *Cep78*^-/-^ mice. Collectively, our findings suggested that CEP78 directly interacts with IFT20 and TTC21A to regulate axonemal and centrosomal functions.

## Discussion

In our study, we found *CEP78* as the causal gene of CRD with male infertility and multiple morphological abnormalities of the sperm flagella using *Cep78^-/-^* mouse. Cep78 forms a trimer with IFT20 and TTC21A (**Figure 8**), which are essential for sperm flagella formation[17, 19]. Cep78 is important for the interaction and stability of the trimer proteins, which regulate cilliogenesis. Absence of CEP78 protein also leaded to CRD and multiple morphological abnormalities of the sperm flagella in human.

**Figure 8.**
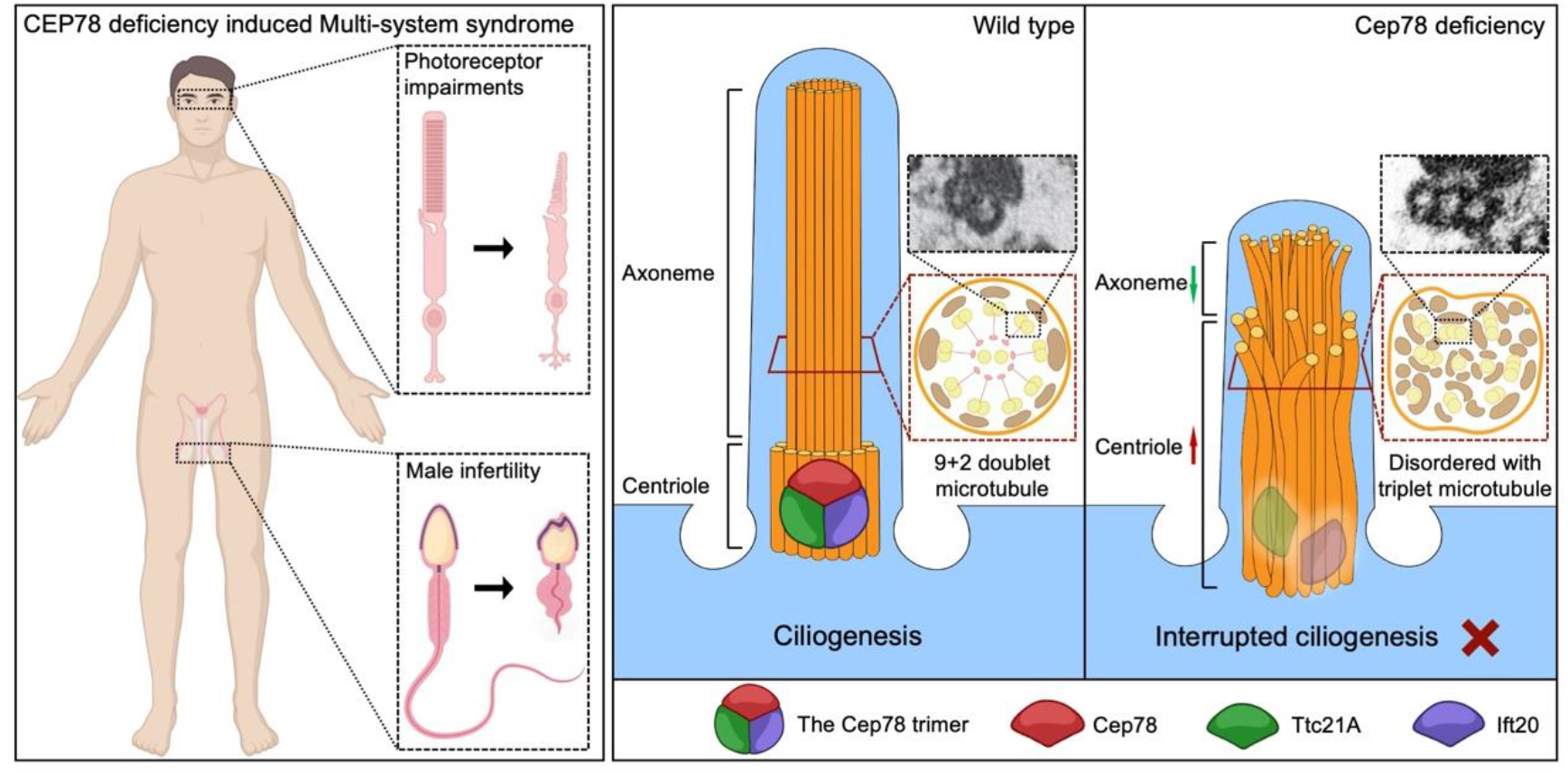
Schematic diagram of pathogenesis induced by Cep78 deficiency. Loss of Cep78 deregulated the expression of and interactions between Ift20 and Ttc21a, two components of the Cep78 trimer. Cep78 deletion causes elongated centrioles, shortened axonemes, abnormal presence of triplet microtubules and disordered 9+2 microtubules.

CEP78 belongs to the centrosomal and ciliary protein family, mutations in other centrosomal genes, such as CEP19, CEP164, CEP250 and CEP290, could disrupt ciliary assembly and generate retinopathy[2, 4, 24-27]. However, there’s no evidence that loss of the above centrosomal proteins can simultaneously cause CRD and male infertility. While in this study, we provide evideces that CEP78 is a causal gene of CRD and male infertility. Absence of Cep78 in human and mouse both present the same phenotypes of CRD, and multiple morphological abnormalities of the sperm flagella.

In *Cep78^-/-^* photoreceptors, we observe shortened connecting cilia, spreading of microtubule doublets in the proximal half, and aberrant organizations of outer segment disks, which is similar to previous findings in mice with disrupted Fam161a protein[28]. Our data provide *in vivo* evidence for CEP78’s role in maintaining structures and functions of microtubule doublets in the connecting cilium of photoreceptors.

Despite of abnormal arrangement of sperm flagella microtubule, we here find that CEP78 disruption could cause abnormal formation of manchette, a transient structure surrounding the elongating spermatid head and is only present during spermatid elongation[22, 29]. During spermiogenesis, manchette and sperm tail axoneme are two microtubular-based platforms that deliver cargo proteins to sperm heads and tails, respectively[29]. How manchette is formed is still not fully elucidated. The majority views hold that manchette microtubules are nucleated at spermatid centrosomes[29, 30], which start from nucleator γ-tubulin of the centrosomal adjunct and end at perinuclear rings[31, 32]. According to our data, other than disorganized manchette structures, elongated centrioles are also observed in *Cep78^-/-^* mice, indicating CEP78’s potential role in organizing manchette microtubules. Taken together, CEP78 dysfunction specifically impairs organization of microtubular-based structures.

Previous study indicated the involvement of CEP78 in regulation of centriole length. We herein also observed stretched centriole length in *Cep78^-/-^* spermatids, and down-regulation of the interacting protein of CEP78, Ift20. CEP78 can recruit the centriole localization of Ift20, whose knock-down causes centriole elongation. Moreover, triplet microtubules, which are specific to proximal centriole in sperm[21], are aberrantly found in principal and middle pieces of sperm flagella from both the male patient with *CEP78* mutation and *Cep78^-/-^* mice, possibly caused by abnormal elongation of the triplet microtubules of centriole. However, such anomaly in both human and mouse sperm has never been reported before.

We found that CEP78 loss leaded to shortening of sperm flagellum, with Cp110 down-regulated in *Cep78*^-/-^ testis (**Supplementary Figure S5**), which is inconsistent with Gonçalves et al.’s findings that cells lacking CEP78 display up-regulated CP110 and reduced cilia frequency with long cilia phenotype[15]. It seems that Cep78 may regulate formation of sperm flagellum by a different mechanism. Our analysis of interacting proteins of Cep78 showed that CEP78 forms a trimer with two other proteins required for sperm flagellar formation, IFT20 and TTC21A. Cep78 regulates the interaction between Ift20 and Ttc21a, and loss of Cep78 leaded to downregulation of the other two components of the trimer. Insufficiency of any component in the trimer disrupts its structure and affects stability of all components, leading to cilia shortening. IFT20, belonging to the IFT family, is essential for assembly and maintenance of cilia and flagella[33]. It is required for opsin trafficking and photoreceptor outer segment development[18], and plays important roles in spermatogenesis and particularly spermiogenesis by transferring cargo proteins for sperm flagella formation[33]. During spermiogenesis, IFT20 is transported along the manchette microtubules[34]. IFT20 is essential to male fertility, spermatogenesis and particularly spermiogenesis by transferring cargo proteins for sperm flagella formation. Multiple sperm flagella abnormalities are observed in *Ift20* mutant mice, including short and kinked tail[19]. TTC21A/IFT139 is also essential for sperm flagellar formation[17]. TTC21A directly interacts with IFT20, and mutations in *TTC21A* gene are reported to cause MMAF in both human and mice[17]. However, more studies are still warranted to better elucidate the biological mechanism of CEP78 in regulating features of spermatozoa and photoreceptors.

CEP78 protein is highly conserved among vertebrate organism with human and mouse CEP78 proteins sharing over 80% similarity. Disease phenotypes and changes of ultra-structures were highly-consistent between human and mice upon CEP78 depletion. Our study also reveals *Cep78^-/-^* mice model as an applicable animal model for therapeutic research of such distinct syndrome.

In conclusion, we identify CEP78 as a causative gene for a type of syndrome involving CRD and male infertility. CEP78 dysfunction induces similar phenotypes and ultra-structure changes in human and mice, including disturbed ciliary structure in photoreceptors, defective sperm flagella structures, and aberrant spermatid head formation. We also found that CEP78 formed a trimer with ciliary proteins IFT20 and TTC21A, and regulated their interaction and stability. The CEP78 trimer is important for the regulation of centriole and cilia length. Our study explains the etiology and animal model of the syndrome of CRD and male infertility, provides basis for molecular diagnosis, and serves as a potential target for gene therapy.

## Materials and Methods

### Ethnical statement

Our study, conformed to the Declaration of Helsinki, was prospectively reviewed and approved by the ethics committee of Ningxia Eye Hospital and Nanjing Medical University. Signed informed consents were obtained from all individuals in the study.

### Mouse breeding

All mice were raised in a specific-pathogen-free animal facility accredited by Association for Assessment and Accreditation of Laboratory Animal Care (AAALAC) in Model Animal Research Center, Nanjing University, China. The facility provided ultraviolet sterilization, a 12-hour light/dark cycle, ad libitum access to water, and standard mouse chow diet. Mice experiments were performed in accordance with approval of the Institutional Animal Care and Use Committee of Nanjing Medical University and with the ARVO Statement for the Use of Animals in Ophthalmic and Vision Research.

### Construction of *Cep78* deficient mice using CRISPR/Cas9

CRISPR/Cas9 technology was applied to generate *Cep78* frameshift mutations in mice using the non-homolog recombination method. Single-guide RNAs (sgRNA) were designed against exons 2 to 11 of *Cep78*. Cas9 mRNA and sgRNAs were synthesized by *in vitro* transcription assay and microinjected into the cytoplasm of single-cell C57BL/6J zygotes. The injected embryos were then transferred into oviducts of pseudopregnant female mice. Generated founder mice and their progenies were genotyped by sequence analyses of the genomic DNA isolated by mice ear clipping to screen for the frameshift mutation in *Cep78*. Genotyping was performed using primer pairs for *Cep78*-null allele with an expected produce size of ∼400 bp and wild type *Cep78* allele with an expected produce size of 517 bp. Primer sequences were listed in **Supplementary Table S5**. Identified founder mice were crossed with wild type C57BL/6J mice to produce F1 progenies. Homozygote *Cep78^-/-^* mice were screened from F2 progenies generated by inbreeding of F1 heterozygote progenies.

### Cell culture and transfection

HEK293T and 3T3 cells were maintained in DMEM medium supplemented with 10% fetal bovine serum (Invitrogen), penicillin (100U/mL) and streptomycin (100g/mL) at 37°C, 5% CO_2_. The open reading frame sequences of mice *Cep78*, *Ift20* and *Ttc21a* were synthesized, amplified and inserted into the pcDNA3.1 plasmid (GenScript, Nanjing, China) with HA/Flag sequences in-frame fused to produce recombinant plasmids Cep78-HA, Ift20-Flag, Ift20-HA and Ttc21a-Flag, respectively. Scramble siRNA, Cep78-siRNA, Ift20-siRNA and Ttc21a-siRNA were purchased from GenePharma (GenePharma, Shanghai, China) with their sequences listed in **Supplementary Table S6**. Transfection assay was conducted using Exfect^®^ transfection reagent (Vazyme, Nanjing, China).

### Immunoblotting

Immunoblotting was conducted according to a previously defined protocol[35, 36]. Mice neural retina and testes were isolated for immunoblotting respectively. Collected tissues were ground in ice-cold protein lysis buffer (Beyotime, Shanghai, China) containing protease inhibitors cocktail (Roche, Basel, Switzerland) for protein extraction. The lysates were separated on 10% sodium dodecyl sulfate-polyacrylamide gel, and then transferred to a polyvinylidene fluoride membrane (Millipore, Billerica, MA, USA). Membranes were then blocked with 5% skim milk at 37°C for one hour, incubated with primary antibodies at 4°C overnight (**Supplementary Table S7**), washed with 1 × tris-buffered saline tween, and probed with corresponding horse radish peroxidase-conjugated secondary antibodies (dilution: 1:10000; ICL Inc., Newberg, Germany) at room temperature for one hour. Blots were developed using the ECL-Western blotting system (Bio-Rad, Hercules, CA, USA).

### ERG

Mice were anesthetized intraperitoneally with a mixture of ketamine (100 mg/kg) and xylazine (10 mg/kg). Pupils were dilated with 1% cyclopentolate-HCL and 2.5% phenylephrine. After dark adaption for 8 hours, ERG was recorded under dim red light using an Espion system (Diagnosys LLC, Lowell, MA, USA) in accordance with recommendations of the International Society for Clinical Electrophysiology of Vision. ERG waves were documented in response to flashes at 0.01, 3.0 and 10.0 cd×s/m^2^.

### SD-OCT

Mice anesthetization and pupil dilation were conducted as above mentioned. Lateral images from nasal retina to temporal retina crossing through the optic nerve were collected using a SD-OCT system (OptoProbe, Burnaby, Canada). Thicknesses of retinal layers were measured by Photoshop software (Adobe, San Jose, CA, USA) in a double-blind manner.

### Immunofluorescence staining

Immunofluorescence staining was performed per a previously described protocol[35, 36]. Mice eyecups and testicular tissues were enucleated after sacrifice, rinsed with 1×phosphate buffer saline (PBS) with connective tissues trimmed, and fixed in 4% paraformaldehyde (PFA) at 4°C overnight. For eyecups, corneas and lens were subsequently removed without disturbing the retina. Posterior eyecups and testicular tissues were then dehydrated in 30% sucrose for 2 hours, embedded in optimal cutting temperature compound, and frozen sectioned at 5 µm. For *in vitro* assay, 3T3 cells were collected at 48 hours post transfection and fixed in 4% PFA at 4°C overnight. Retinal, testicular and cellular sections were then blocked in 2% normal goat serum and permeabilized with 0.3% Triton X-100 at room temperature for 1 hour. Those sections were further incubated with primary antibodies (**Supplementary Table S7**) at 4°C overnight, and corresponding fluorescence-conjugated secondary antibodies (dilution: 1:1000; Invitrogen, Carlsbad, CA, USA) at room temperature for 1 hour. Nuclei were counterstained with 4’, 6-diamidino-2-phenylindole (DAPI; Sigma, St. Louis, MO, USA). Retinal images were collected with a Leica TCS SP5 confocal system (Leica), and testicular and cellular images were taken by an LSM 800 confocal microscope (Carl Zeiss, Jena, Germany).

### TEM and SEM assays

Mice eyecups, mice semen cells collected from shredded unilateral caudal epididymis, mice testicular tissues, and human ejaculated sperm were subjected to TEM assay, respectively. Tissues were fixed in 2.5% glutaradehyde at 4°C overnight immediately after enucleation. Eyecups were dissected as described for TEM, and only the remaining posterior eyecups were used for the following experiments. Samples were then post-fixed in 1% osmic acid at 4°C for an hour, stained in aqueous 3% uranyl acetate for 2 hours, dehydrated with ascending concentrations of acetone, embedded in epoxy resin, cut into ultra-thin slides, and stained in 0.3% lead citrate. Ultrastructure of mice retina and human spermatozoa was visualized using a JEM-1010 electron microscope (JEOL, Tokyo, Japan). Ultrastructure of mice spermatids cells were observed with a Philips CM100 electron microscope (Philips, Amsterdam, North-Holland). Ultrastructure of testicular tissues was visualized using a FEI Tecnai G2 Spirit Bio TWIN electron microscope (Thermo, Waltham, MA, USA).

For SEM assay, mice sperms collected from shredded unilateral caudal epididymis and human ejaculated sperm were used. Human and mice spermatozoa were immersed in 2.5% glutaraldehyde at 4°C overnight, fixed in 1% osmic acid supplemented with 1.5% K_3_[Fe(CN)_3_] at 4°C for an hour, steeped in 1% thiocarbohydrazide for an hour, post-fixed in 1% osmic acid at 4°C for an hour, and soaked in 2% uranyl acetate solution at 4°C overnight. The treated samples were then progressively dehydrated with an ethanol and isoamyl acetate gradient on ice, and dried with a CO2 critical-point dryer (Eiko HCP-2, Hitachi Ltd., Toyko, Japan). The specimens were subsequently mounted on aluminum stubs, sputtercoated by an ionic sprayer meter (Eiko E-1020, Hitachi Ltd.), and visualized using a FEI Nova NanoSEM 450 scanning electron microscope (Thermo).

### CASA and sperm morphological study

Mice sperms, collected from shredded unilateral caudal epididymis, were maintained in 500 µL modified human tubal fluid (Irvine Scientific, Santa Ana, CA, USA) supplemented with 10% fetal bovine serum (Gibco, Grand Island, NY, USA) at 37 °C for 7 min. Sperm concentration, motility, and progressive motility of semen samples were then assessed by CASA (IVOS II, Hamilton Thorne Inc., Beverly, MA, USA).

Human ejaculated semen sample was collected by masturbation after 7 days of sexual abstinence. Semen volume was measured with test tube. After liquefaction at 37 °C for 30 min, semen sample was further subjected to CASA (BEION S3-3, BEION, Beijing, China) for measurement of sperm concentration, motility and progressive motility per the 5^th^ edition of World Health Organization (WHO) guidelines.

Human and mice semen samples were stained on slides using the H&E staining method for sperm morphological analyses. At least 150 spermatozoa were analyzed for each group. Percentages of spermatozoa with normal morphology, abnormal head, abnormal neck and abnormal flagella were calculated per the WHO guidelines. Abnormalities of sperm flagella were further classified into 7 categories, including short flagella, absent flagella, coiled flagella, multi flagella, cytoplasm remain, irregular caliber, and angulation. Noteworthy, one spermatozoon was sorted into only one group based on its major flagellar abnormality.

### Histological H&E staining

For histological H&E staining, mice epididymal tissues and sperms collected from shredded unilateral caudal epididymis were rinsed with 1×PBS, fixed in 4% PFA at 4°C overnight, dehydrated with ascending concentrations of ethanol, embedded in paraffin, and sectioned into 5 µm slides. Slides were then deparaffinized in xylene, rehydrated with decreasing concentrations of ethanol, and stained with hematoxylin and eosin for histological observation. Images were taken with a Zeiss Axio Skop plus2 microscope (Carl Zeiss).

### PAS staining

PAS staining was used for spermatogenic staging and evaluation of the development of seminiferous tubules. Fresh mice testicular tissues were fixed with modified Davidson’s fluid (mDF) fixation fluid at room temperature for 48 hours, embedded with paraffin, and sliced into sections. Slides were then dewaxed with xylene at 37°C for 30 min, rehydrated by descending concentrations of ethanol, dyed with periodic acid, Schiff (Solarbio, Beijing, China) and hematoxylin in sequence, dehydrated with ascending concentrations of ethanol, and sealed with neutral balsam for observation. Images were taken with a Zeiss Axio Skop plus2 microscope (Carl Zeiss).

### Sample preparation, IP, and MS analyses

Testicular tissues of *Cep78^+/-^* and *Cep78^-/-^* mice were lysed with Pierce^TM^ IP lysis buffer (Thermo) supplemented with 1% protease inhibitor cocktail 100X (Selleck Chemicals, Houston, TX, USA), respectively. Lysates were revolved for 1 hour at 4°C and centrifuged at 40000g for 1 hour. Supernatants were precleared with 20 μL of protein A/G magnetic beads (Millipore) at 4°C for 1 hour. IP was performed using the Pierce^TM^ co-IP kit (Thermo). Eluted proteins were then subjected to SDS-PAGE, silver stain, immunoblotting and MS. For MS, silver stained gel lanes were carefully cut into small pieces and were then processed for MS analysis. Briefly, after trypsin digestion overnight, peptides were desalted using stage tips, re-suspended in 0.1% formic acid (v/v) and subjected to LC-MS[37–40]. Data of IP-MS was submitted to Dryad, Dataset, https://doi.org/10.5061/dryad.6djh9w12z.

For quantitative mass spectrometry (MS) on elongating spermatids lysates of *Cep78^+/-^* and *Cep78^-/-^* mice, lysed samples were digested overnight at 37 ◦C with trypsin and subjected to the TMT labeling. For MS analyses, each fraction was analyzed using an Orbitrap Fusion Lumos mass spectrometer (Thermo Finnigan, San Jose, CA) coupled with Easy-nLC 1200 (Thermo Finnigan, San Jose, CA), and was separated by a High PH reverse phase saperation microcapillary column (ACQUITY BEH C18 Column, 1.7 μm, 300μm X 150 mm) at a flow rate of 4μL/min in a 128-min linear gradient (3% buffer B for 14 min, 8% buffer B for 1 min, 29% buffer B for 71 min, 41% buffer B for 12 min, 100% buffer B for 9 min, 3% buffer B for 21 min) (solvent A: 20mM Ammonium formate, PH=10; solvent B: 100% ACN, 0.1% FA). Sample were collected at a rate of one tube per minute, a total of 30 components. Easy1200 and fusion Lumos were used for series identification. Data of quantitative mass spectrometry (MS) on elongating spermatids lysates was submitted to Dryad, Dataset, https://doi.org/10.5061/dryad.stqjq2c4p

### co-IP assay

HEK293T cells were collected at 48 hours post transfection, and lysed with Pierce^TM^ IP lysis buffer (Thermo) supplemented with 1% protease inhibitor cocktail 100X (Selleck Chemicals). Lysates was revolved for 30 minutes at 4°C and centrifuged at 16000g for 30 minutes. Supernatants were incubated with 30 μL of Pierce^TM^ anti-HA magnetic beads (Thermo) or anti-DYKDDDDK IP resin (GenScript) at 4°C overnight. IP was performed using the Pierce^TM^ co-IP kit (Thermo), and eluted proteins were further subjected to immunoblotting.

### RNA extraction and Q-PCR

3T3 cells were harvested at 48 hours post transfection for RNA extraction. Total RNA was isolated from lysates of transfected 3T3 cells using TRIzol reagent (Invitrogen). RNA concentration and quality were determined with Nano-Drop ND-1000 spectrophotometer (Nano-Drop Technologies, Wilmington, DE, USA). cDNA was generated with a PrimeScript RT Kit (Takara, Otsu, Shiga, Japan). Q-PCR was conducted to detect RNA amounts using FastStart Universal SYBR Green Master (ROX; Roche, Basel, Switzerland) with StepOne Plus Real-Time PCR System (Applied Biosystems, Darmstadt, Germany). Primer sequences were listed in **Supplementary Table S5**.

### Statistical analysis

GraphPad Prism (version 8.0; GraphPad Software, San Diego, CA, USA) was applied for statistical analyses. Student’s t test was utilized for comparisons between two different groups. One-way analysis of variance (ANOVA) followed by Dunnet’s multiple comparison test was applied for comparisons among three or more groups. We presented data as mean ± standard error of the mean, and considered *p* < 0.05 as statistically significant. All experiments were conducted in both biological and technical triplicates with data averaged.

## Acknowledgements

We thank all participants for their sample donations. This work was supported by the National Key R&D Program (2021YFC2700200 to X.G); National Natural Science Foundation of China (82020108006, 81730025 to C.Z, 81971439, 81771641 to X.G, 82070974 to X.C, 82060183 to X.S); Shanghai Outstanding Academic Leaders (2017BR013 to C.Z); and Six Talent Peaks Project in Jiangsu Province (YY-019 to X.G). The funders have no role in study design, data collection and analysis, decision to publish, or preparation of the manuscript.

## Author Contributions

X.C., X.G. and C.Z. conceived and designed the study. T.Z. and Y.Z. conducted experiments. T.Z and X.C. interpreted the data and drafted the manuscript. X.G. and C.Z. revised the manuscript. X.S., X.Z., Y.C., Y.S., and Y.Q. coordinated and analyzed the data. All the authors contributed to, read, and approved the final manuscript.

## References

1. Gill, J.S., et al., Progressive cone and cone-rod dystrophies: clinical features, molecular genetics and prospects for therapy. Br J Ophthalmol, 2019.

2. Kubota, D., et al., CEP250 mutations associated with mild cone-rod dystrophy and sensorineural hearing loss in a Japanese family. Ophthalmic Genet, 2018. 39(4): p. 500–507.

3. Charbel Issa, P., et al., Olfactory Dysfunction in Patients With CNGB1-Associated Retinitis Pigmentosa. JAMA Ophthalmol, 2018. 136(7): p. 761–769.

4. Yildiz Bolukbasi, E., et al., Homozygous mutation in CEP19, a gene mutated in morbid obesity, in Bardet-Biedl syndrome with predominant postaxial polydactyly. J Med Genet, 2018. 55(3): p. 189–197.

5. Brunk, K., et al., Cep78 is a new centriolar protein involved in Plk4-induced centriole overduplication. J Cell Sci, 2016. 129(14): p. 2713–8.

6. Hossain, D., et al., Cep78 controls centrosome homeostasis by inhibiting EDD-DYRK2-DDB1(Vpr)(BP). EMBO Rep, 2017. 18(4): p. 632–644.

7. Azimzadeh, J., et al., Centrosome loss in the evolution of planarians. Science, 2012. 335(6067): p. 461–3.

8. Fu, Q., et al., CEP78 is mutated in a distinct type of Usher syndrome. J Med Genet, 2017. 54(3): p. 190–195.

9. Nikopoulos, K., et al., Mutations in CEP78 Cause Cone-Rod Dystrophy and Hearing Loss Associated with Primary-Cilia Defects. Am J Hum Genet, 2016. 99(3): p. 770–776.

10. Namburi, P., et al., Bi-allelic Truncating Mutations in CEP78, Encoding Centrosomal Protein 78, Cause Cone-Rod Degeneration with Sensorineural Hearing Loss. Am J Hum Genet, 2016. 99(3): p. 777–784.

11. Sanchis-Juan, A., et al., Complex structural variants in Mendelian disorders: identification and breakpoint resolution using short- and long-read genome sequencing. Genome Med, 2018. 10(1): p. 95.

12. Ascari, G., et al., Functional characterization of the first missense variant in CEP78, a founder allele associated with cone-rod dystrophy, hearing loss, and reduced male fertility. Hum Mutat, 2020.

13. Zhang, M., et al., Low expression of centrosomal protein 78 (CEP78) is associated with poor prognosis of colorectal cancer patients. Chin J Cancer, 2016. 35(1): p. 62.

14. Nesslinger, N.J., et al., Standard treatments induce antigen-specific immune responses in prostate cancer. Clin Cancer Res, 2007. 13(5): p. 1493–502.

15. Goncalves, A.B., et al., CEP78 functions downstream of CEP350 to control biogenesis of primary cilia by negatively regulating CP110 levels. Elife, 2021. 10.

16. Bedoni, N., et al., Mutations in the polyglutamylase gene TTLL5, expressed in photoreceptor cells and spermatozoa, are associated with cone-rod degeneration and reduced male fertility. Hum Mol Genet, 2016. 25(20): p. 4546–4555.

17. Liu, W., et al., Bi-allelic Mutations in TTC21A Induce Asthenoteratospermia in Humans and Mice. Am J Hum Genet, 2019. 104(4): p. 738–748.

18. Keady, B.T., Y.Z. Le, and G.J. Pazour, IFT20 is required for opsin trafficking and photoreceptor outer segment development. Mol Biol Cell, 2011. 22(7): p. 921–30.

19. Zhang, Z., et al., Intraflagellar transport protein IFT20 is essential for male fertility and spermiogenesis in mice. Mol Biol Cell, 2016.

20. Zhu, X., et al., Mouse cone arrestin expression pattern: light induced translocation in cone photoreceptors. Mol Vis, 2002. 8: p. 462–71.

21. Avidor-Reiss, T. and E.L. Fishman, It takes two (centrioles) to tango. Reproduction, 2019. 157(2): p. R33–R51.

22. Kierszenbaum, A.L. and L.L. Tres, The acrosome-acroplaxome-manchette complex and the shaping of the spermatid head. Arch Histol Cytol, 2004. 67(4): p. 271–84.

23. Fu, Q., et al., CEP78 is mutated in a distinct type of Usher syndrome. J Med Genet, 2017. 54(3): p. 190–195.

24. Chaki, M., et al., Exome capture reveals ZNF423 and CEP164 mutations, linking renal ciliopathies to DNA damage response signaling. Cell, 2012. 150(3): p. 533–48.

25. Valente, E.M., et al., Mutations in CEP290, which encodes a centrosomal protein, cause pleiotropic forms of Joubert syndrome. Nat Genet, 2006. 38(6): p. 623–5.

26. Baala, L., et al., Pleiotropic effects of CEP290 (NPHP6) mutations extend to Meckel syndrome. Am J Hum Genet, 2007. 81(1): p. 170–9.

27. den Hollander, A.I., et al., Mutations in the CEP290 (NPHP6) gene are a frequent cause of Leber congenital amaurosis. Am J Hum Genet, 2006. 79(3): p. 556–61.

28. Karlstetter, M., et al., Disruption of the retinitis pigmentosa 28 gene Fam161a in mice affects photoreceptor ciliary structure and leads to progressive retinal degeneration. Hum Mol Genet, 2014. 23(19): p. 5197–210.

29. Lehti, M.S. and A. Sironen, Formation and function of the manchette and flagellum during spermatogenesis. Reproduction, 2016. 151(4): p. R43–54.

30. Kierszenbaum, A.L., et al., Ran, a GTP-binding protein involved in nucleocytoplasmic transport and microtubule nucleation, relocates from the manchette to the centrosome region during rat spermiogenesis. Mol Reprod Dev, 2002. 63(1): p. 131–40.

31. Akhmanova, A., et al., The microtubule plus-end-tracking protein CLIP-170 associates with the spermatid manchette and is essential for spermatogenesis. Genes Dev, 2005. 19(20): p. 2501–15.

32. Kierszenbaum, A.L., E. Rivkin, and L.L. Tres, Cytoskeletal track selection during cargo transport in spermatids is relevant to male fertility. Spermatogenesis, 2011. 1(3): p. 221–230.

33. Zhang, Z., et al., Intraflagellar transport protein IFT20 is essential for male fertility and spermiogenesis in mice. Mol Biol Cell, 2016. 27(23): p. 3705–16.

34. Kazarian, E., et al., SPAG17 Is Required for Male Germ Cell Differentiation and Fertility. Int J Mol Sci, 2018. 19(4).

35. Jiang, C., et al., MicroRNA-184 promotes differentiation of the retinal pigment epithelium by targeting the AKT2/mTOR signaling pathway. Oncotarget, 2016. 7(32): p. 52340–52353.

36. Liu, Y., et al., SPP2 Mutations Cause Autosomal Dominant Retinitis Pigmentosa. Sci Rep, 2015. 5: p. 14867.

37. Guo, X., et al., Proteomic analysis of male 4C germ cell proteins involved in mouse meiosis. Proteomics, 2011. 11(2): p. 298–308.

38. Guo, X., et al., Proteomic analysis of proteins involved in spermiogenesis in mouse. J Proteome Res, 2010. 9(3): p. 1246–56.

39. Hu, Z., et al., A genome-wide association study identifies two risk loci for congenital heart malformations in Han Chinese populations. Nat Genet, 2013. 45(7): p. 818–21.

40. Fan, Y., et al., Phosphoproteomic Analysis of Neonatal Regenerative Myocardium Revealed Important Roles of Checkpoint Kinase 1 via Activating Mammalian Target of Rapamycin C1/Ribosomal Protein S6 Kinase b-1 Pathway. Circulation, 2020. 141(19): p. 1554–1569.

